# Loss of Adaptive DNA Breaks in Alzheimer’s Disease Brains

**DOI:** 10.1101/2023.12.11.566423

**Authors:** Xiaoyu Zhang, Mohammad Haeri, Russell H. Swerdlow, Ning Wang

## Abstract

**Background:** DNA breaks accumulate in Alzheimer’s disease (AD) brains. While their role as true genomic lesions is recognized, DNA breaks also support cognitive function by facilitating the expression of activity-dependent immediate early genes (IEGs). This process involves TOP2B, a DNA topoisomerase that catalyzes the formation of DNA double-strand breaks (DSBs).

**Objective:** To characterize how AD impacts adaptive DNA breaks at nervous system genes.

**Methods:** We leveraged the ability of DNA single- and double-strand breaks to activate poly(ADP-ribose) polymerases (PARPs) that conjugate poly(ADP-ribose) (PAR) to adjacent proteins. To characterize the genomic sites harboring DNA breaks in AD brains, nuclei extracted from 3 AD and 3 non-demented (ND) autopsy brains (frontal cortex, all male donors, age 78 to 91 years of age) were analyzed through CUT&RUN in which we targeted PAR with subsequent DNA sequencing.

**Results:** Although the AD brains contained 19.9 times more PAR peaks than the ND brains, PAR peaks at nervous system genes were profoundly lost in AD brains, and the expression of these genes was downregulated. This result is consistent with our previous CUT&RUN targeting γH2AX, which marks DNA double-strand breaks (DSBs). In addition, TOP2B expression was significantly decreased in the AD brains.

**Conclusion:** Although AD brains contain a net increase in DNA breaks, adaptive DNA breaks at nervous system genes are lost in AD brains. This could potentially reflect diminished TOP2B expression and contribute to impaired neuron function and cognition in AD patients.

## Introduction

Alzheimer’s disease (AD) is a progressive neurodegenerative disorder that results in cognitive impairment and memory loss [1]. AD causes the majority of dementia cases worldwide, and has increasingly become a global challenge as the population ages [2]. Although brain extracellular amyloid plaques and intraneuronal neurofibrillary tangles are considered pathological hallmarks of AD and are well characterized [3], the mechanistic cause of the cognitive symptoms in AD remains unclear.

Mounting evidence reveals damage to the genome, in the form of DNA breaks, accumulates in AD brain neurons [4–12]. Post-mitotic, long-lived neurons are particularly vulnerable to DNA damage as their genome integrity must be carefully maintained. Thus, DNA breaks likely contribute to AD neurodysfunction and neurodegeneration.

In neurons, though, DNA breaks also play a physiological role. In response to external stimuli, neurons rapidly form recurrent DNA double-strand breaks (DSBs) at genomic loci near immediate early genes (IEGs), which facilitate their transcription [13, 14]. IEG expression subsequently initiate profound adaptations in neural circuit morphology and connectivity through the activation of late-response genes; this process is critical to experience-driven memory formation [15]. DNA topoisomerase II-β (TOP2B) mediates this process by catalyzing these adaptive DSBs [16, 17]. In addition, high levels of single strand breaks (SSBs) frequently arise at neuronal enhancers [18, 19]. By relieving topological constraints, DNA breaks induce gene transcription, which in turn modify neuronal circuits and plasticity.

We recently used the CUT&RUN approach that targets γH2AX, a marker for DSBs, to compare DSB distributions in brains from AD and age- and sex-matched non-demented (ND) individuals [20]. In the study we now report, we extended that finding by CUT&RUN that targets ADP-ribose (PAR), a marker for both SSBs and DSBs [21], to map DNA break sites. Our results collectively show that DNA breaks are detected at genes whose gene ontology (GO) functions are enriched for nervous system-related genes in ND samples. Interestingly and importantly, despite an overall increase in DNA breaks in AD samples, DNA breaks at these genes are lost in AD brains. DNA breaks are instead formed at ectopic genomic sites with functions related to protein ubiquitination and catabolic processes.

## MATERIALS AND METHODS

### Human samples

Autopsy brain (frontal cortex) tissues were obtained from the University of Kansas Alzheimer’s Disease Research Center (KU ADRC) Neuropathology Core. For information on the human samples used in this study (AD, n = 3 males; controls, n = 3 males), please refer to **Table 1**.

**Table 1.** Details of AD and ND samples. CAA, cerebral amyloid angiopathy. LB, Lewy body.

| ID | Sex | Age | Diagnosis | Tissue |
| --- | --- | --- | --- | --- |
| ADC3 | M | 82 | ND | FCx, Superior B1 |
| ADC5 | M | 89 | ND | FCx, Superior B1 |
| ADC7 | M | 86 | AD/CAA/TDP43+ | FCx, Superior B1 |
| ADC11 | M | 87 | AD/LB | FCx, Superior B1 |
| ADC13 | M | 78 | AD/CAA | FCx, Superior B1 |
| ADC14 | M | 91 | ND | FCx, Superior B1 |

### Immunohistochemistry

For immunohistochemistry, FFPE histological sections were dewaxed, rehydrated in an ethanol series (95%, 85%, and 70%) and then subjected to microwave antigen retrieval in 0.01 M citrate (pH 6.0; 0.05% Tween-20) and methanol/H_2_O_2_ treatment. After blocking with 5% goat serum, the slides were sequentially incubated with anti-poly-ADP-ribose binding reagent (Millipore-Sigma, MABE1031) at a 1:1,000 dilution and HRP-labeled secondary antibodies. A NovaRed kit (Vector, SK-4800) was used for visualization.

### Preparation of nuclei from human brain specimens

Nuclei were extracted from frozen human brain tissues by using Nuclei EZ Prep (Millipore-Sigma, NUC-101) following the manufacturer’s manual. The detailed procedures are described in [20]. Briefly, after thawing on ice, human brain tissues were minced by blades and homogenized in a Dounce homogenizer. The homogenate was then filtered through a 70-µm mesh strainer and centrifuged at 500 x g for 5 min at 4 °C. After removing the supernatant, the nuclei were resuspended in 1.5 ml of Nuclei EZ Lysis Buffer and incubated for another 5 min on ice. The nuclei were centrifuged at 500 × g for 5 min at 4 °C. After removing the supernatant, the nuclei were washed and then filtered through a 40-µm mesh strainer. Intact nuclei were counted after counterstaining with Trypan blue in a standard cell counter.

### CUT&RUN

For CUT&RUN, 500,000 nuclei were washed 1x with CUT&RUN wash buffer (20 mM HEPES, pH 7.5, 150 mM NaCl, 0.5 mM spermidine), bound to activated ConA beads, permeabilized in wash buffer (wash buffer + 0.002% digitonin), incubated with anti-poly-ADP-ribose binding reagent, washed in wash buffer, incubated with pA-MN (EpiCypher 15-1016), and washed in wash buffer. Following the final wash, the cells were washed with ice-cold low-salt wash buffer and digested using MNase digestion buffer for 25 min on ice. Solubilized chromatin was released using an isosmotic stop buffer and collected using a PCR cleanup kit column.

CUT&RUN Library Prep was performed using an Illumina NovaSeq 6000 Sequencing System at the University of Kansas Medical Center Genomics Core (Kansas City, KS). Fragmented input and immunoprecipitated chromatin (5 ng) were used to initiate the TruSeq ChIP Sample Prep Kit library preparation protocol with modifications for CUT&RUN sample input (Illumina Cat# IP-202-1012). The fragmented chromatin underwent end repair and 3’ adenylation prior to Illumina indexed adapter ligation. No gel size selection of the ligation product was performed. Ten cycles of PCR amplification with a modified extension time of 10 seconds were performed using Illumina adaptor-specific priming with final library purification using KAPA Pure magnetic bead purification (KAPA Cat# KK8002).

Library validation was performed using a DNA 1000 Assay Kit (Agilent Technologies 5067-1504) on an Agilent TapeStation 4200. The concentration of each library was determined by qPCR using a Roche LightCycler 96 using FastStart Essential DNA Green Master Mix (Roche 06402712001) and KAPA Library Quant (Illumina) DNA Standards 1-6 (KAPA Biosystems KK4903). The libraries were pooled based on equal molar amounts to 1.85 nM for multiplexed sequencing.

The pooled libraries were denatured with 0.2 N NaOH (0.04 N final concentration) and neutralized with 400 mM Tris-HCl pH 8.0. Dilution of the pooled libraries to 370 pM was performed in the sample tube on the instrument, after which onboard clonal clustering of the patterned flow cell was performed using a NovaSeq 6000 S1 Reagent Kit v1.5 (200 cycle) (Illumina 20028318). A 2x101 cycle sequencing profile with dual index reads was completed using the following sequence profile: Read 1 – 101 cycles x Index Read 1 – 6 cycles x Index Read 2 – 0 cycles x Read 2 – 101 cycles. Following collection, the sequence data were converted from the .bcl file format to the .fastq file format using bcl2fastq software and demultiplexed into individual sequences for data distribution using a secure FTP site or Illumina BaseSpace for further downstream analysis.

### CUT&RUN data processing and analysis

TrimGalore was used to trim the raw .fastq files to remove adaptors. The trimmed .fastq files were then mapped to the hg19 genome utilizing Bowtie2. The same procedure was run to align the .fastq files to a masked *Saccharomyces cerevisiae* v3 (sacCer3) genome for spike-in control DNA, which was also downloaded from the University of California Santa Cruz (UCSC) (https://genome.ucsc.edu/). Sambamba was then used to remove duplicates. For IGV visualization, deepTools was used with the “bamCoverage” function to generate normalized CPM .bw files. For peak calling, the recently developed SEACR was utilized and run in “relaxed’ mode to produce peak files, as the BED files used were already normalized to the number of yeast spike-in reads. DeepTools was further applied for heatmap visualization with the functions “computeMatrix” and “plotHeatmap”. The “dba.peakset” function of the Diffbind R package was further applied to identify overlapping peaks on the basis of the bound peaks. The links of CUT&RUN peaks and their related genes were established with the “annotatePeaks.pl” function in Homer. Motif enrichment analyses were performed using the “findMotifsGenome.pl” function in Homer, leading to known enrichment results and *de novo* enrichment results, and the latter were chosen in this study.

## Results

### Loss of DSB sites in nervous system genes in AD

In our previous CUT&RUN targeting γH2AX [20], in addition to predominantly observing an increase in the number of γH2AX peaks in AD brains, there were also 12,167 γH2AX peaks that were downregulated. These downregulated γH2AX peaks were distributed in 3’ and 5’ untranslated regions (UTRs), exons, intergenic regions, introns, and promoters (**Fig. 1A and B**). Interestingly, in gene ontology (GO) analysis, γH2AX peaks in the intronic regions and intergenic regions were enriched for nervous system functions, e.g., dendrite development (*SHANK3*, *CAMK2A*), synapse organization (*ARC*, *MECP2*) (**Fig. 1C**). These data suggest that while DNA DSBs are formed in the intronic and intergenic regions of nervous system function genes in ND brains, presumably to facilitate their expression, these DSBs are lost in AD brains.

**Fig. 1.**
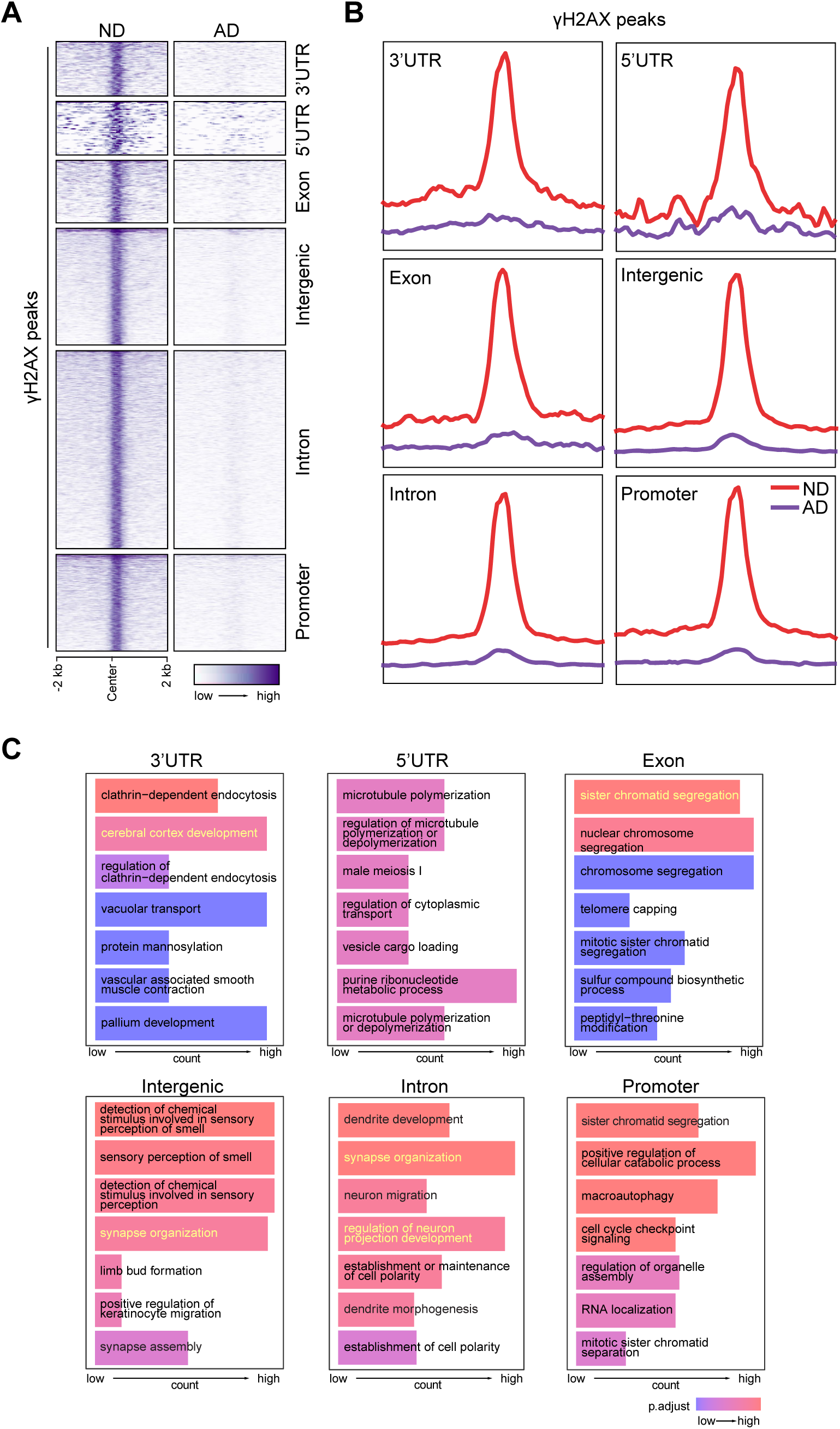
Loss of CUT&RUN γH2AX peaks in AD brain. **(A)** Heatmaps showing distribution of γH2AX peaks at 3’UTR, 5’UTR, exon, intergenic region, intron, and promoter. **(B)** Distribution of γH2AX peaks in a ±2 kb window at 3’UTR, 5’UTR, exon, intergenic region, intron, and promoter. **(C)** GO analysis for PAR peaks at 3’UTR, 5’UTR, exon, intergenic region, intron, and promoter.

### Mapping DNA break sites using CUT&RUN targeting PAR

Our prior CUT&RUN measured DSBs indirectly through adjacent γH2AX [20]. However, γH2AX can be found up to 50 kb away from both sides of DSBs in euchromatin [22]. Thus, to confirm our results, we performed CUT&RUN targeting ADP-ribose (PAR) by using an anti-PAR binding reagent. Various types of DNA damage, including DSBs, SSBs, and single-strand gaps, activate PAR polymerases (PARPs). PARP activity signals the presence of DNA damage by modifying adjacent proteins with PAR. Thus, CUT&RUN for PAR can be used to detect DNA breaks and map their genomic location [18].

We obtained autopsy brain tissue from the frontal cortex of postmortem human ND and AD patients through the University of Kansas Alzheimer’s Disease Research Center (KU ADRC) Neuropathology Core. The age, sex, and diagnostic information is shown in **Table 1**. First, we examined PAR-associated DNA damage in these samples. Immunohistochemistry (IHC) for PAR using histological sections from these patients showed a few cells positive for PAR in ND samples; in contrast, a significant increase in the number of cells positive for PAR was observed in AD samples, especially in neurons (74.4 ± 10.5 in AD samples vs 16.8 ± 6.9 in ND samples, *P* < 0.05) (**Fig. 2A**).

**Fig. 2.**
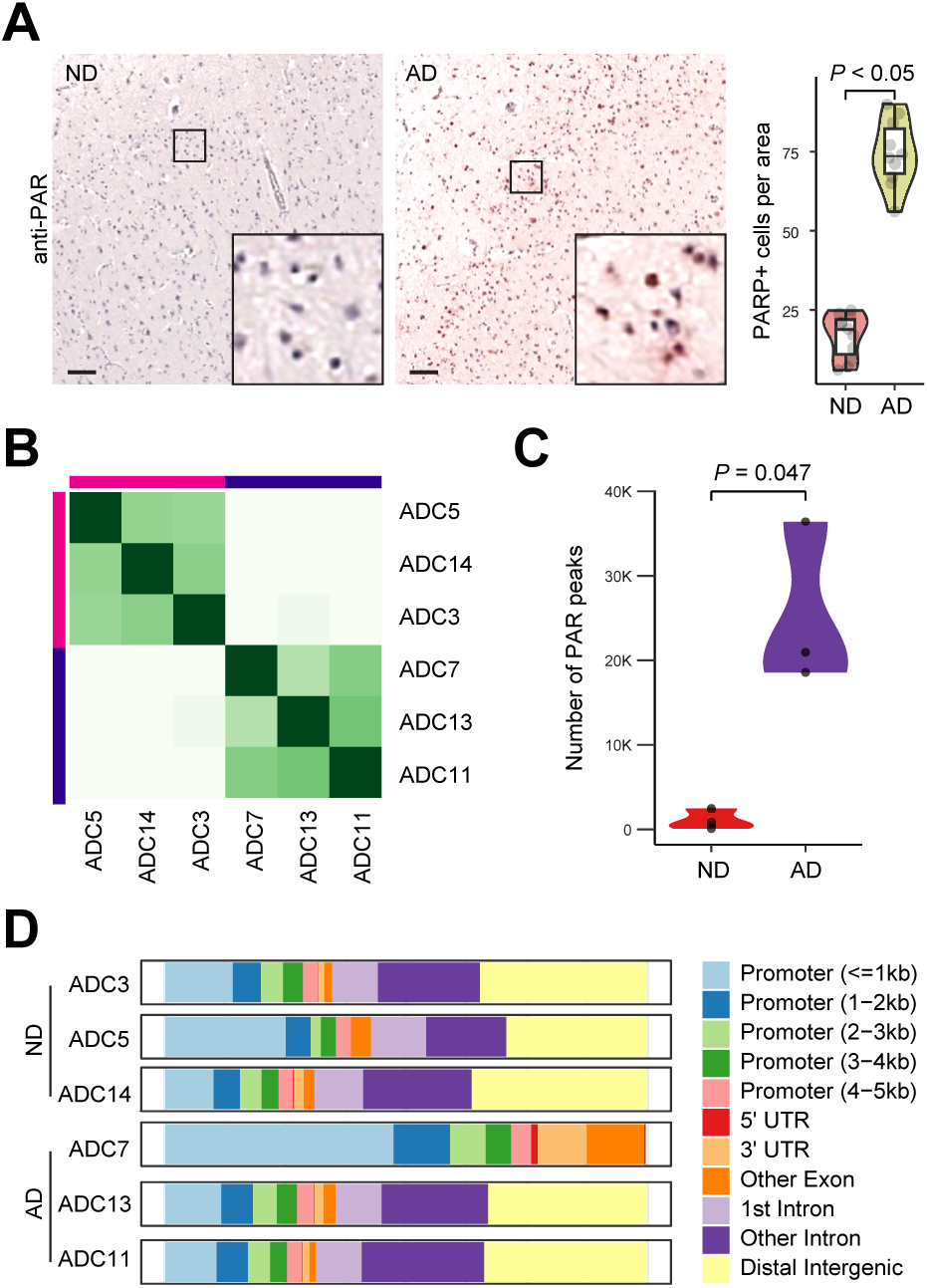
CUT&RUN for PAR analysis in AD brain. **(A)** Increased PAR staining by IHC in AD samples. Scale bar, 100 μm. PAR-positive cells were quantified from multiple view areas in 3 ND and 3 AD samples. Data represent mean ± SD. **(B)** Diffbind analysis show clustering of CUT&RUN for PAR in AD and ND samples. **(C)** Elevated PAR peaks (DNA breaks) in AD vs ND samples. **(D)** Distribution of differential PAR peaks in genome.

Following CUT&RUN, differential binding analysis of CUT&RUN and hierarchical clustering revealed that all 3 ND samples (ADC3, ADC5 and ADC14) were clustered together, while all 3 AD samples (ADC7, ADC11, and ADC13) were clustered together, indicating a high degree of conservation in the DNA break profile between ND and AD samples (**Fig. 2B**). There was a 19.9-fold increase in the number of PAR peaks detected in AD versus ND samples (**Fig. 2C**). Thus, consistent with previous reports, our CUT&RUN for PAR data shows an accumulation of DNA damage in AD. Furthermore, distribution analysis shows that many PAR peaks are in the intergenic and intronic regions in both ND and AD samples (**Fig. 2D**), except ADC7, which shows a profound loss of CUT&RUN PAR peaks in the intergenic and intronic regions (γH2AX peaks are not lost in ADC7 in these genomic regions [20]).

### Altered genomic distribution of PAR peaks in AD brains

CUT&RUN for PAR shows strong central peaks, which presumably indicate DNA break sites (**Fig. 3A, B, D, and E**). GO analysis shows that these DNA break sites upregulated in AD occur at genes that encode regulators of proteasome-mediated ubiquitin process (e.g., *UBE2J2*, *DVL1*, *PARK7*), small molecule catabolic process (e.g., *ENO1*, *PGD*, *ALDH4A1*), and histone modification (e.g., *NOC2L*, *PRDM2*, *PADI2*), etc. (**Fig. 3C**). Kyoto Encyclopedia of Genes and Genomes (KEGG) analysis identified protein processing in endoplasmic reticulum (e.g., *FBXO2*, *FBXO6*), carbon metabolism (e.g, *SDHB*, *PKLR*), etc. (**Fig. 3C**).

**Fig. 3.**
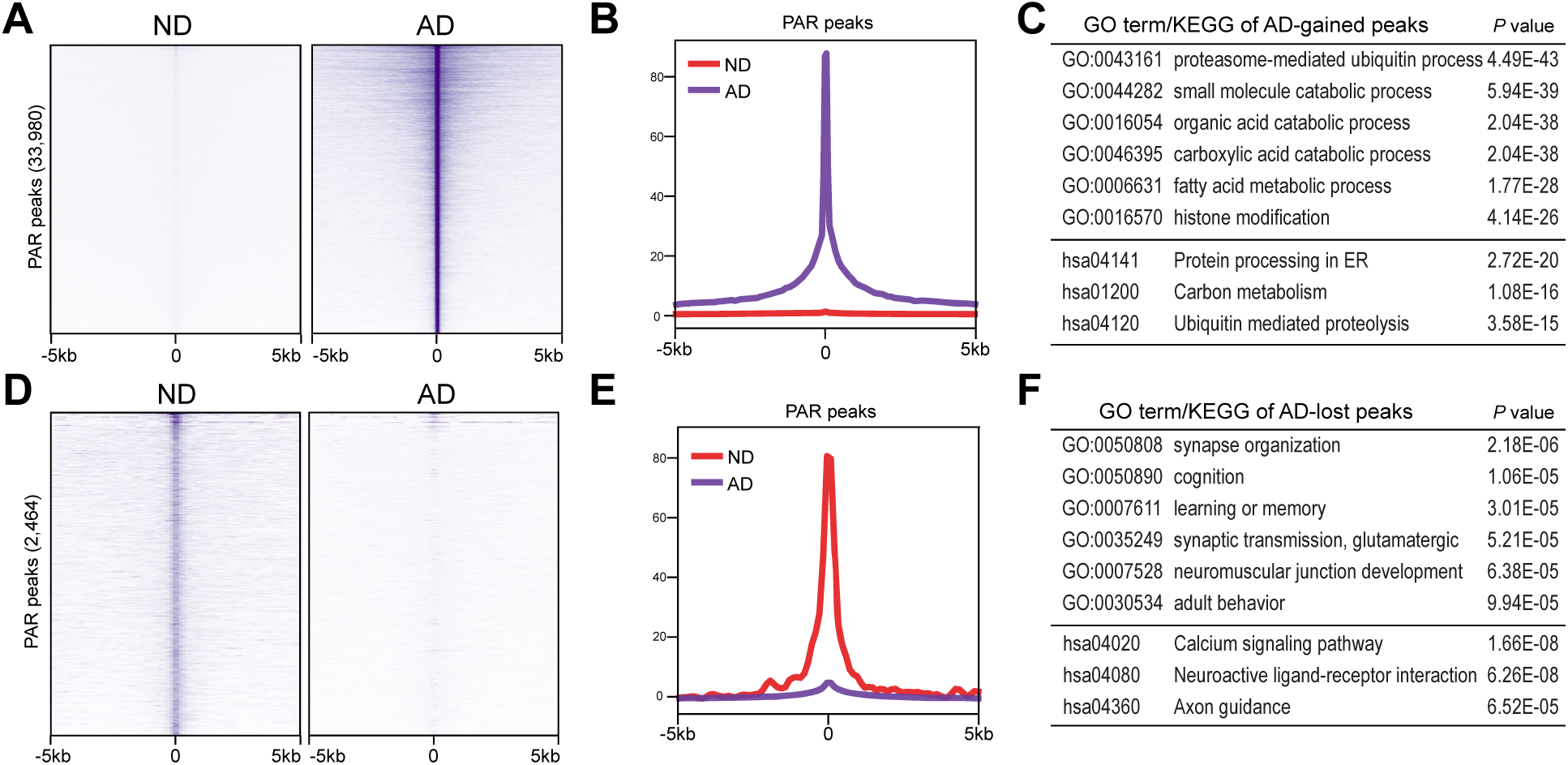
PAR peaks in ND and AD samples. **(A, D)** Heatmaps showing distribution of AD-enriched **(A)** and ND-enriched **(D)** PAR peaks. **(B, E)** Distribution of ±5 kb window of AD-enriched **(B)** and ND-enriched **(E)** PAR peaks. **(C, F)** GO and KEGG analysis of AD-enriched **(C)** and ND-enriched **(F)** PAR peaks.

Interestingly, GO analysis of DNA break sites downregulated in AD revealed genes that encode regulators of synapse organization (e.g., *NFIA*, *ARHGAP22*, *ANK3*), cognition (e.g., *PRKCZ*, *FOXO6*, *SORCS3*), glutamatergic synaptic transmission (e.g., *GRIK3*, *LRRK2*, *SYT1*), etc. (**Fig. 3F**). KEGG analysis identified calcium signaling pathway (e.g., *PDE1C*, *CAMK2D*, *PLCE1*), neuroactive ligand-receptor interaction (e.g., *PRL*, *CHRNA7*, *CHRM5*), Axon guidance (*SEMA6A*, *CAMK2D*, *SLIT2*), etc. (**Fig. 3F**).

Together, these data suggest DNA breaks that form in genes mediating nervous system function are lost in AD. Instead, AD-associated DNA breaks are strongly accumulated at ectopic genomic sites associated with novel functions, such as ubiquitin and catabolic processes.

### PAR peaks upregulated in AD samples

We examined the enrichment profiles of PAR peaks in ND and AD samples, which showed distribution around the 3’-UTR, 5’UTR, exons, and promoters (**Fig. 4A and B**). Interestingly, our data show that CUT&RUN PAR peaks at these genomic regions are enriched for different functions in functional pathway identification analysis (**Fig. 4C**). While PAR peaks at the promoters and exons are uniquely enriched for pathways related to cadherin binding (e.g., *CDH11*, *CTNND1*) and ATP hydrolysis activity (e.g., *ASCC2*, *FBX01*), PAR peaks at 3’-UTR and 5’-UTRs encode pathways related to SNARE binding (e.g., *UVRAG*, *VAMP7*) and ligase activity (e.g., *ASCL4*, *SCL27A2*). Moreover, functional term identification analysis (**Fig. 3D**) showed that PAR peaks at promoters and exons are uniquely enriched for many functions, such as protein processing in endoplasmic reticulum (e.g., *CUL3*, *CTAGE4*) and regulation of actin cytoskeleton (e.g., *ROCK1*, *WASHC5*). Only 248 PAR peaks were upregulated in the intron and intergenic regions and functional pathway identification analysis shows that they are enriched for functions, such as cellular response to nutrient levels (e.g., *BCL2*, *LAMP2*) and cellular response to extracellular stimulus (e.g., *ATG7*, *GSDMD*) (data now shown).

**Fig. 4.**
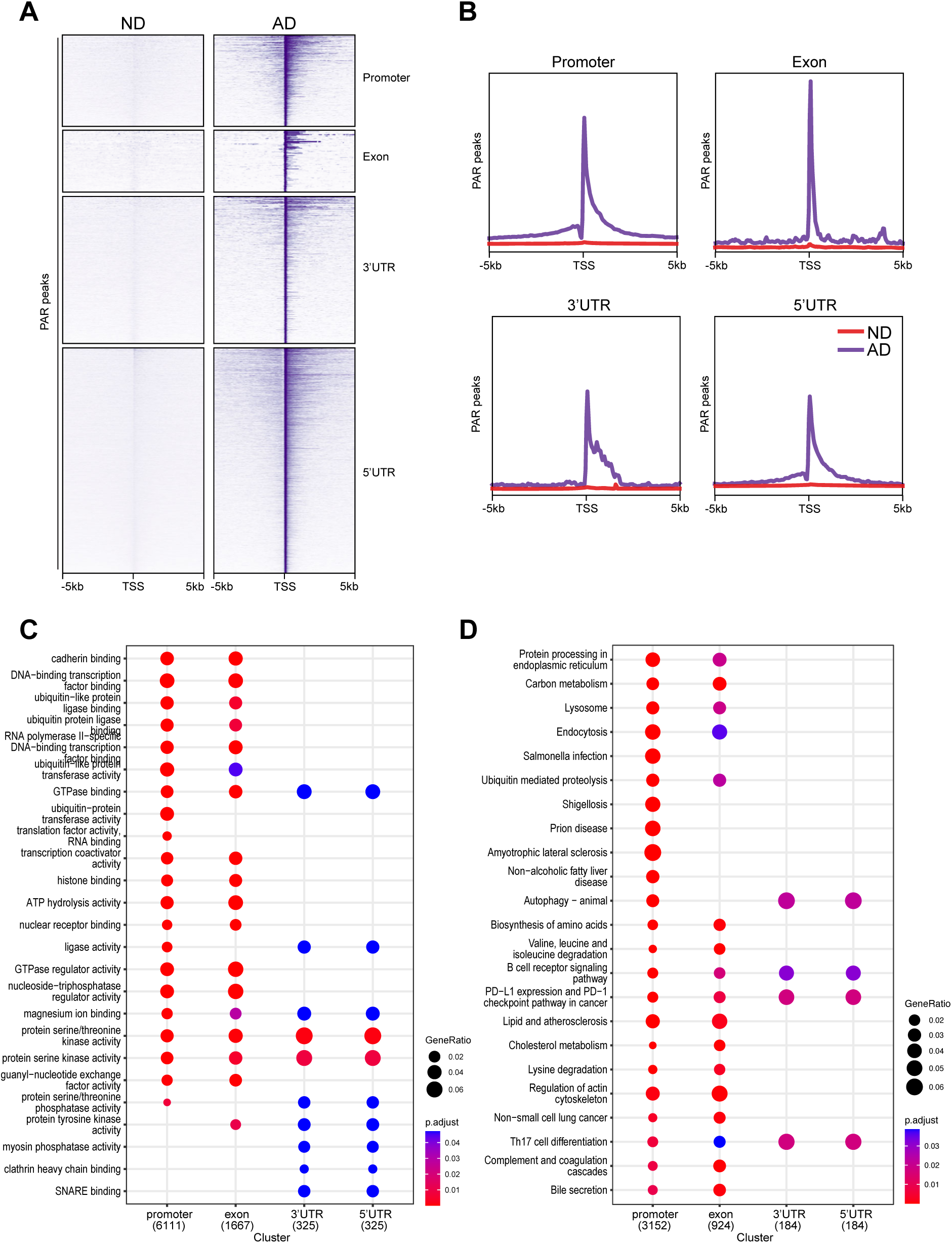
Genomic distribution of PAR peaks upregulated in AD samples with functional analysis. **(A)** Heatmaps showing distribution of PAR peaks at promoter, intron, exon, 3’UTR and intergenic. **(B)** Distribution of PAR peaks in a ±2 kb window of γH2AX binding sites at promoter, intron, exon, 3’UTR and intergenic. **(C)** GO analysis for PAR peaks at promoter, exon, 3’UTR and 5’UTR. **(D)** KEGG analysis for PAR peaks at promoter, exon, 3’UTR and 5’UTR.

### PAR peaks downregulated in AD samples

We further examined the enrichment profiles of CUT&RUN PAR peaks in ND and AD samples, which showed distribution around the promoters, introns, intergenic regions, and exons (**Fig. 5A and B**). Interestingly, functional pathway identification analysis demonstrates that PAR peaks at the promoters are enriched for functions, such as protein localization to nucleus (e.g., *CDK1*, *CTNNA1*), at the exons are enriched for functions, such as inorganic cation import across plasma membrane (e.g., *ABCC9*, *ATP1A1*), and at intergenic regions are enriched for functions, such as regulation of protein catabolic process (e.g., *ABCA2*, *ADAM8*) (**Fig. 5C**). Importantly, similar to the intronic gH2AX peaks downregulated in AD (**Fig. 1**), PAR peaks at introns downregulated in AD are also enriched for nervous system-related functions (e.g., *ADCY1*, *CHD7*) (**Fig. 5C**).

**Fig. 5.**
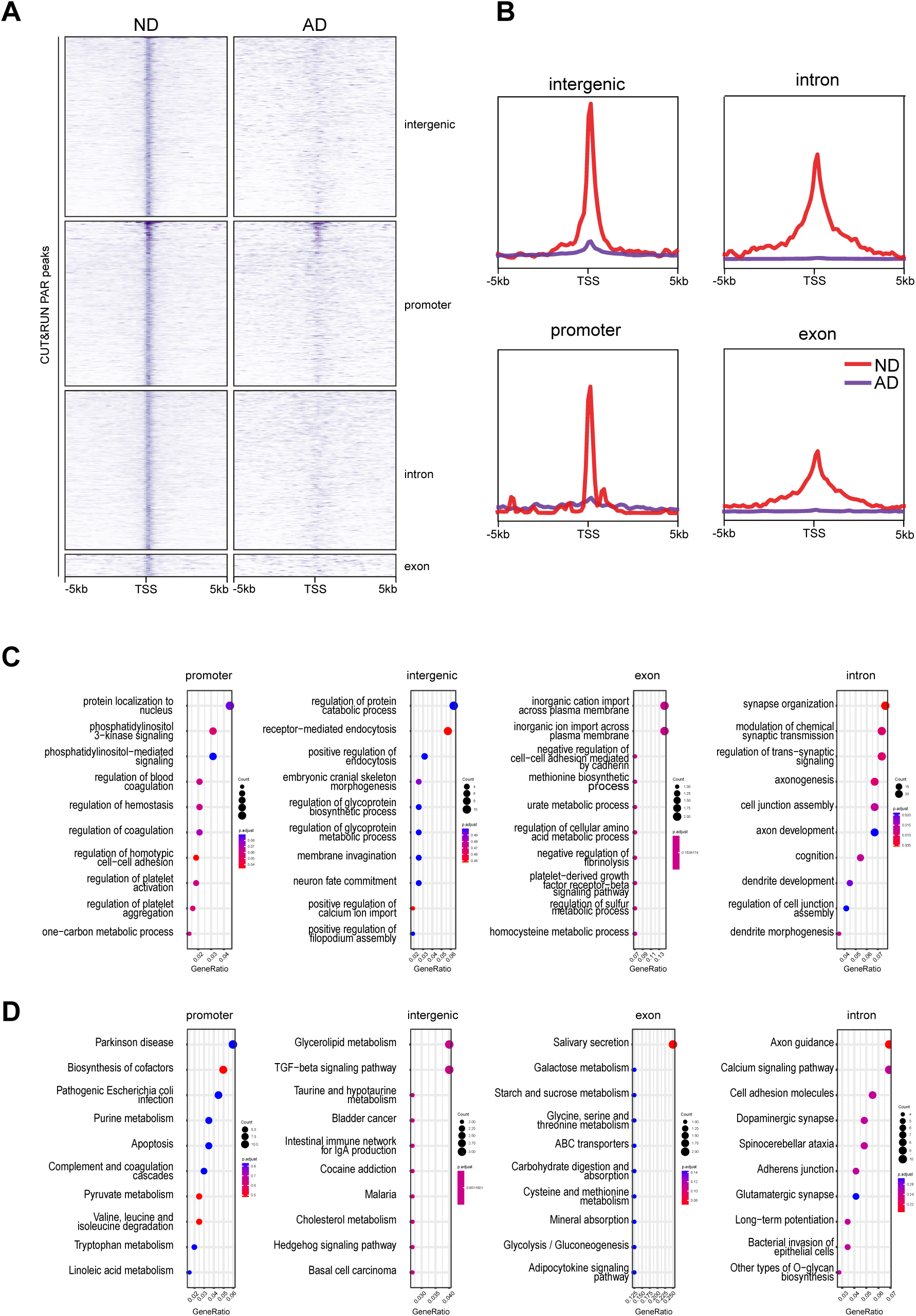
Genomic distribution of PAR peaks downregulated in AD samples with functional analysis. **(A)** Heatmaps showing distribution of PAR peaks at promoter, intron, intergenic, and exon. **(B)** Distribution of PAR peaks in a ±2 kb window of γH2AX binding sites at promoter, intron, intergenic, and exon. **(C)** GO analysis for PAR peaks at promoter, intron, intergenic, and exon. **(D)** KEGG analysis for PAR peaks at promoter, intron, intergenic, and exon.

### Differential PAR peaks in AD samples

We performed differential peak analysis on DNA breaks located near transcription start sites (TSSs) (**Fig. 6A**) and used Hypergeometric Optimization of Motif EnRichment (HOMER) reference to examine sequences related to transcription factor (TF)-binding sites. Our analysis showed that while the binding motifs for the CEBP and CREB TFs were among the most enriched in ND-associated peaks (**Fig. 6B**), the binding motifs for the Olig2 and RARα TFs were among the most enriched in AD-associated peaks (**Fig. 6C**).

**Fig. 6.**
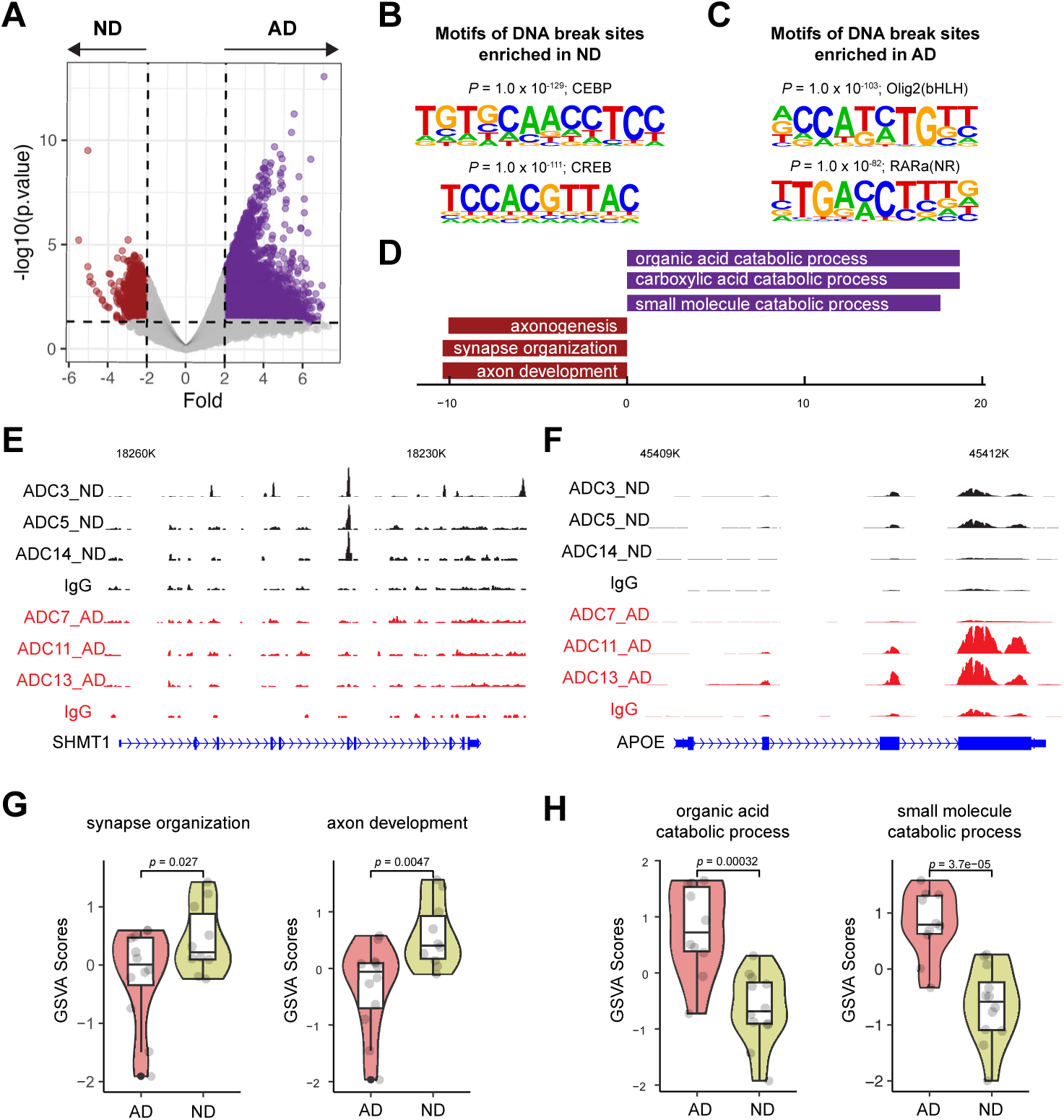
Differential PAR peak analysis and correlation with gene expression. **(A)** Differentially enriched PAR peaks between AD and ND. Number of differential peaks and peaks with >2-fold change is labeled. **(B)** Sequences and significance of enrichment of DNA motifs at PAR in ND samples. **(C)** Sequences and significance of enrichment of DNA motifs at PAR in AD samples. **(D)** GO analysis of biological processes for AD and ND enriched peaks. **(E)** Representative image showing identified peaks binding in the *SHMT1* gene locus from AD and ND. **(F)** Representative image showing identified peaks binding in the *APP* gene locus from AD and ND. **(G)** Violin plots showing the expression of representative ND-associated gene signature levels. **(H)** Violin plots showing the expression of representative AD-associated gene signature levels.

After linking peaks to respective genes using the Genomic Regions Enrichment of Annotations Tool (GREAT) [23], we identified 8,326 genes with upregulated- and 638 genes with downregulated PAR occupancy in AD samples, suggesting that these genes have increased and decreased numbers of DNA breaks in AD samples, respectively. Next, we analyzed their functions. GO analysis showed that the genes with upregulated PAR occupancy in AD samples were enriched for functions such as the organic acid catabolic process; in contrast, genes with downregulated PAR occupancy in AD samples were enriched for nervous system-related functions such as axon development, synapse organization, etc. (**Fig. 6D**). For example, *SHMT1* encodes a folate-dependent enzyme that mediates neuronal and cognitive function by regulating *de novo* thymidylate biosynthesis [24]. PAR peaks were detected at the *SHMT1* gene in ND but not AD samples (**Fig. 6E**), suggesting that DNA breaks are formed at the *SHMT1* gene in ND samples but not in AD samples. Conversely, PAR peaks were detected in the *APOE* gene in some AD samples but not in any of the ND samples (**Fig. 6F**).

To identify the potential impact of the observed altered DNA breaks on gene expression, we examined the gene expression profile in an RNA-seq dataset of 10 ND samples and 12 AD samples [25]. We found that genes associated with nervous system function-related GO terms (“synapse organization”, “axon development”, etc.) was significantly downregulated in AD samples (**Fig. 6G**). In contrast, the expression of genes with GO terms (“organic acid catabolic process”, “small molecule catabolic process”) with AD-associated DNA breaks was upregulated in AD samples (**Fig. 6H**). These data suggest that AD-related loss and gain of DNA breaks in the genes bearing these GO functions result in their down- and upregulated gene expression, respectively.

### Downregulation of TOP2B expression in AD brains

DNA topoisomerases catalyze DNA break formation. To test whether altered DNA breaks results from dysregulated DNA topoisomerase expression in AD, we initially examined the expression of DNA topoisomerases (TOP1, TOP2A, TOP2B, TOP3A, and TOP3B) using the RNA-seq dataset [25]. Our data show that only the expression of TOP2B and TOP3B significantly change in AD. Specifically, TOP2B expression is downregulated in AD (**Fig. 7A**), while TOP3B is aberrantly upregulated in AD (data not shown).

**Fig. 7.**
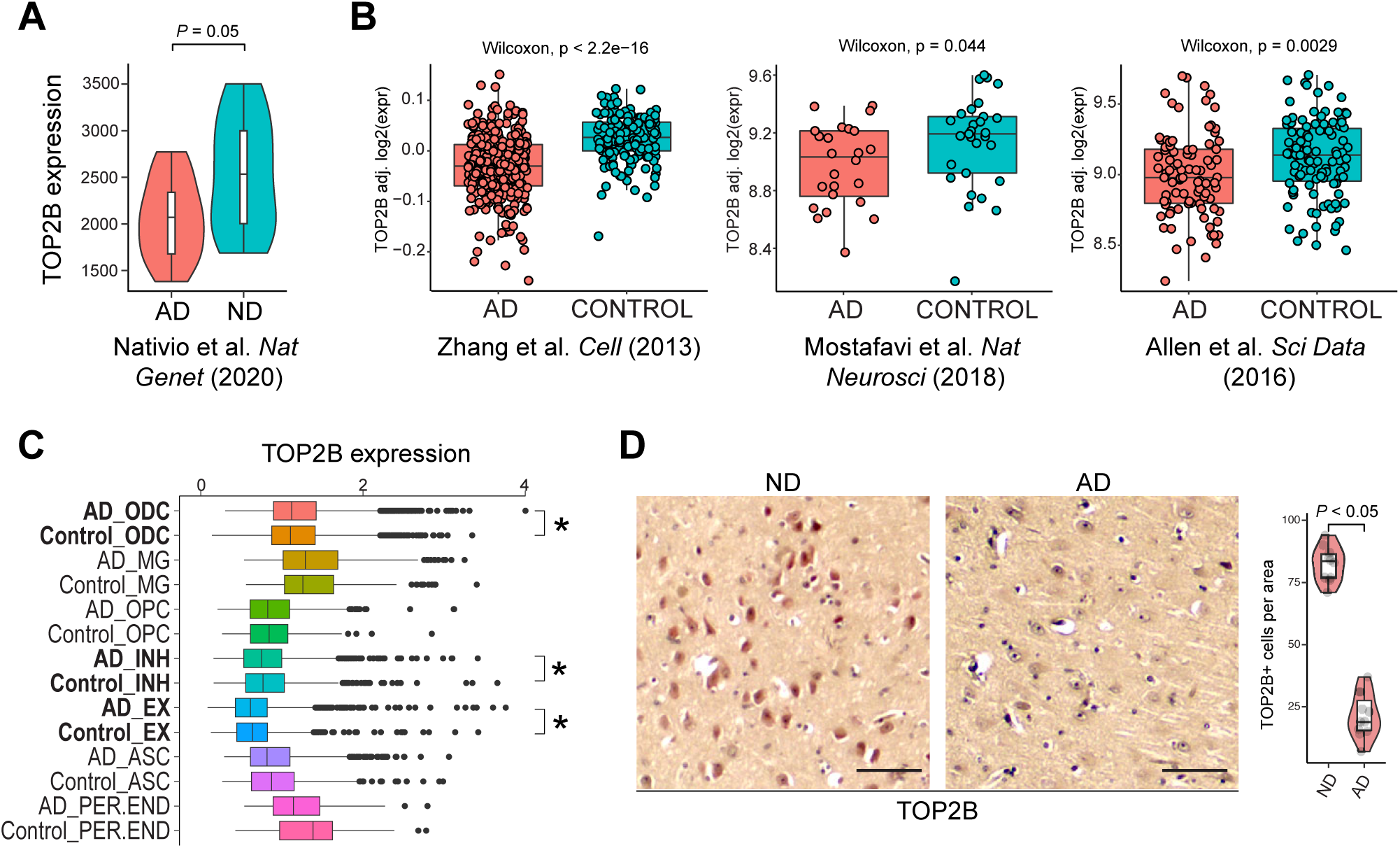
TOP2B expression in ND and AD brains. **(A)** TOP2B expression is reduced in AD in the indicated RNA-seq dataset. **(B)** TOP2B expression is reduced in AD in three published RNA-seq datasets. **(C)** scRNA-seq: TOP2B is downregulated in INH and EX neurons but upregulated in ODC neurons. *, p < 0.05. **(D)** Reduced TOP2B IHC staining in AD vs ND brains. Scale bar, 50 μm. TOP2B-positive cells were quantified from multiple view areas in 3 ND and 3 AD samples. Data represent mean ± SD.

TOP2B plays a critical role in producing DSBs to facilitate IEG expression, which mediates synaptic plasticity following neuronal activity [16, 17]. Thus, downregulated TOP2B expression is consistent with a loss of DNA breaks near genes associated with nervous system function genes. We further examined TOP2B expression in three published large-scale RNA-seq studies using ND and AD samples [26–28]. In all three datasets, *TOP2B* expression was significantly downregulated in AD brains (**Fig. 7B**). By investigating a single cell RNA sequencing (scRNA-seq) dataset [29], we found that *TOP2B* expression was significantly downregulated in AD brain inhibitory (INH) and excitatory (EX) neurons, and upregulated in oligodendrocytes (ODC) (**Fig. 7C**). To examine TOP2B expression at the protein level we conducted IHC staining, which showed that there is a significantly decreased number of cells expressing TOP2B in AD brains (21.7 ± 9.3 in AD samples vs 82.4 ± 7.0 in ND samples, *P* < 0.05) (**Fig. 7D**). Together, these data are consistent with the possibility that loss of DNA breaks in genes critical nervous system function may reflect TOP2B expression in AD brains.

## DISCUSSION

Although it has been long known that DNA breaks are accumulated in AD neurons, their functional consequences are unclear. Our study shows there is significant accumulation of DNA breaks at ectopic genomic loci with functions related to catabolic processes, and that the expression of those genes are upregulated. This data suggests that DNA breaks may dysregulate neuronal metabolism. In addition, our study shows a loss of DNA breaks whose GO functions are enriched related nervous system function. This may result from or contribute to impaired neuronal function in AD. As DNA breaks are known to alter the morphology and connectivity of neural circuits by regulating gene transcription [30], this could perturb neuron function and health.

There was a significant overlap between DNA breaks mapped through CUT&RUN targeted to γH2AX or PAR (**Fig. 8**). While γH2AX is considered as a specific marker of DSBs, PAR labels both SSBs and DSBs (and single-strand gaps). However, ataxia telangiectasia and Rad3-related protein (ATR) can phosphorylate H2AX in response to SSBs [31, 32]. Thus, in our CUT&RUN targeting γH2AX, some SSBs may also present as γH2AX peaks.

**Fig. 8.**
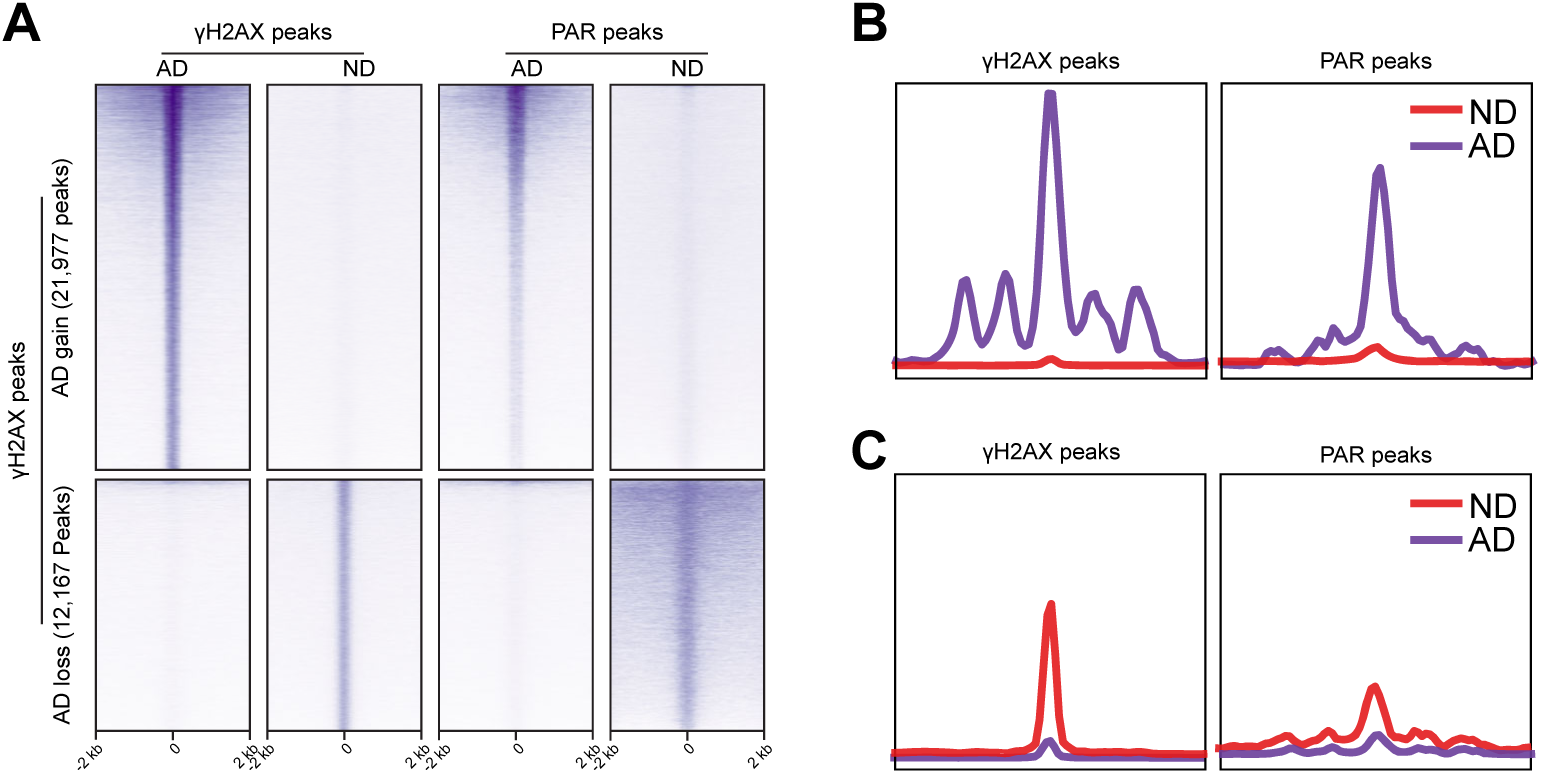
Comparison of differential γH2AX and PAR peaks in ND and AD samples. **(A)** Heatmaps showing distribution of γH2AX and PAR peaks. **(B)** Distribution of γH2AX and PAR peaks in a ±2 kb window upregulated in AD. **(C)** Distribution of γH2AX and PAR peaks in a ±2 kb window downregulated in AD.

An increase in AD brain PAR indicates elevated PARP activity in AD that results from the accumulation of DNA damage. Interestingly, PARPs for DNA repair are major NAD+- and ATP-consuming enzymes in cells, and their sustained PARP activation may deplete NAD+- and ATP level and cause mitochondrial dysfunction and impaired energy metabolism. On the other side, mitochondrial dysfunction and impaired energy metabolism are hallmarks of AD [33–35], which in turn could impact DNA repair and exacerbate the DNA damage in AD neurons.

Limitations of our current study include small sample size and the use of only frontal cortex samples. Future studies that increase the sample size and include hippocampal samples could provide a more comprehensive understanding of alterations of DNA break patterns in AD. In addition, past studies have demonstrated that Top2β mediates activity-dependent neuronal DNA breaks in mouse brains [16]. Although our results demonstrate that a loss of both functional DNA breaks and TOP2B expression in human AD brains, our study has yet been able to establish a causal relationship between these two events. Interestingly, we have conducted a preliminary study of TOP2B target genes using a ChIP-seq dataset that computes TOP2B binding sites (GSE141528) [36]. The data shows that ND-associated DNA breaks (lost in AD samples) are DNA targets of TOP2B (**Fig. 9A**). Among these genes bound by TOP2B include *ARC*, a classical IEG that encodes a master regulator of synaptic function and memory formation [37]. The regulation of *ARC* by TOP2B is further supported by that *ARC* expression is downregulated in TOP2B-deficient SH-SY5Ycells (GSE142383) (**Fig. 9B**) [38]. Our data shows that the ARC gene lost DSBs in human AD samples (**Fig. 9C**). We further examined ARC expression in a large-scale RNA-seq studies, in which ARC expression is significantly down-regulated in AD brains [28] (**Fig. 9D**). By examining a scRNA-seq dataset [29], we found that ARC was downregulated EX neurons (and astrocytes) in AD brains (**Fig. 9E**). However, a past study in mice show that the *Arc* gene does not show activity-induced DNA breaks and its expression is not affected by *Top2β* knockdown [16]. Thus, future studies are needed to formally establish whether reduced TOP2B expression leads to the loss of functioning DNA breaks around genes that are enrich for nervous system-related functions in AD brains. This would be important to understand the causes that underlie impaired neuron and cognitive function in AD patients.

**Fig. 9.**
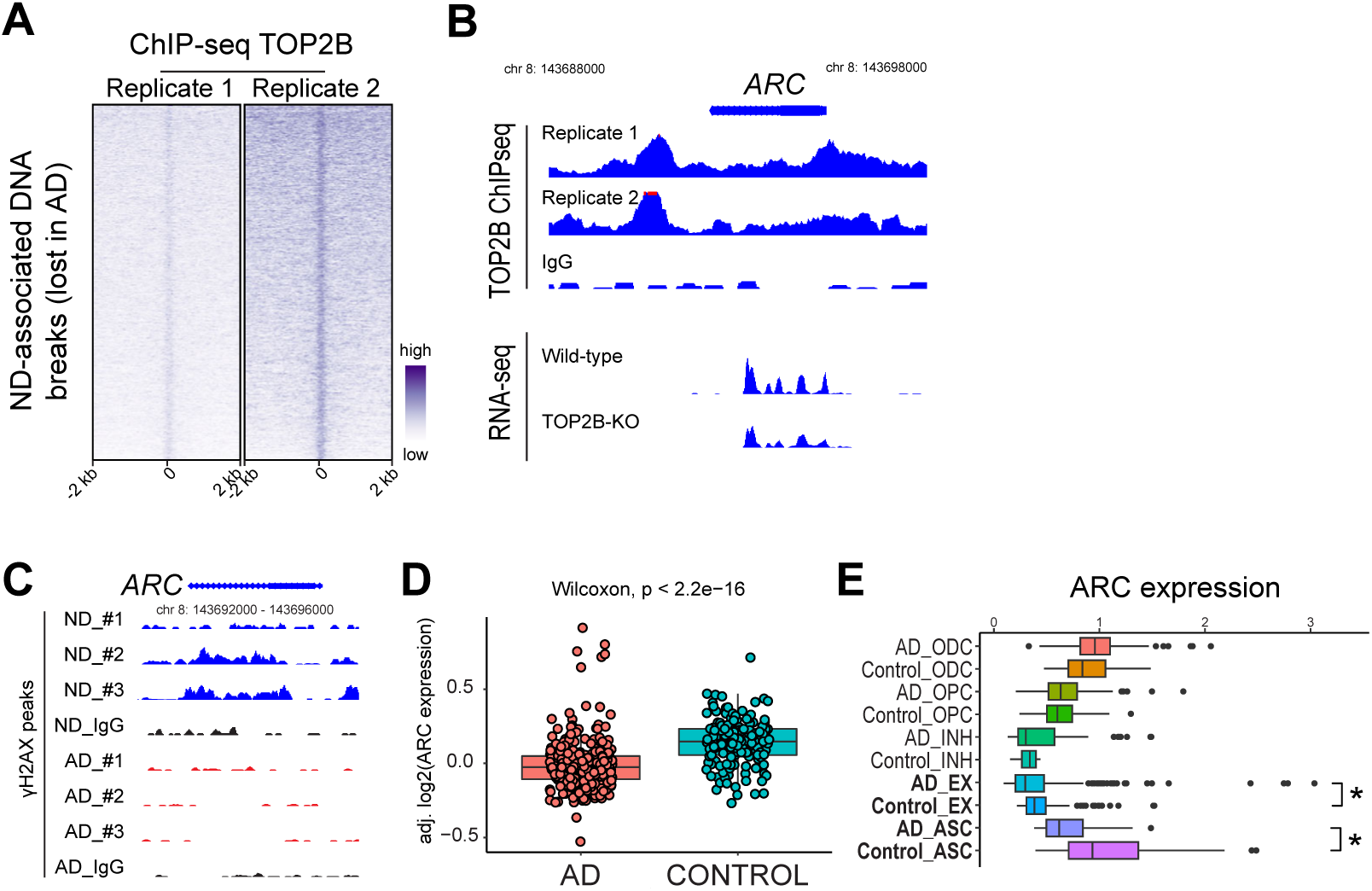
Functioning ND DNA breaks (lost in AD) are TOP2B target genes. **(A)** Heatmap of TOP2B ChIP-seq. **(B)** Genome track for the *ARC* gene in TOP2B ChIP-seq and RNA-seq in WT and TOP2B-deficient SH-SY5Y cells. **(C)** Genome track for the *ARC* gene in ND and AD samples in γH2AX CUT&RUN. **(D)** *ARC* expression is decreased in AD in a published RNA-seq dataset. **(E)** scRNA-seq: ARC expression is downregulated in AD EX neurons. *, p < 0.05.

## ACKNOWLEDGEMENTS

We thank Cynthia Shaddy-Gouvion at KUMC Histopathology Core for preparing frozen autopsied tissues and FFPE sections.

## FUNDING

This study was supported by R01-HD103888, KU ADRC P30 AG072973, the KU School of Medicine, and the Landon Center on Aging (N.W.). This project was supported by an Institutional Development Award (IDeA) from the National Institute of General Medical Sciences of the National Institutes of Health under grant number P20 GM103418 (X.Z.). Research reported in this publication was supported by the KUMC Research Institute, Inc (X.Z.). The content is solely the responsibility of the authors and does not necessarily represent the official views of these funders. Kansas Intellectual and Developmental Disabilities Research Center (NIH U54 HD 090216), the Molecular Regulation of Cell Development and Differentiation – COBRE (P30 GM122731-03) - the NIH S10 High-End Instrumentation Grant (NIH S10OD021743) and the Frontiers CTSA grant (UL1TR002366) at the University of Kansas Medical Center, Kansas City, KS 66160.

## CONFLICT OF INTEREST

The authors have no conflict of interest to report.

## DATA AVAILABILITY

The dataset used in the current study has been uploaded and will be made publicly available at NCBI.

## Notes

### Competing Interest Statement

The authors have declared no competing interest.

## REFERENCE

[1] Apostolova LG (2016) Alzheimer disease. Continuum: Lifelong Learning in Neurology 22, 419.

[2] Cummings JL (2004) Alzheimer’s disease. New England Journal of Medicine 351, 56–67.

[3] Chen X-Q, Mobley WC (2019) Alzheimer disease pathogenesis: insights from molecular and cellular biology studies of oligomeric Aβ and tau species. Frontiers in Neuroscience 13, 659.

[4] Adamec E, Vonsattel JP, Nixon RA (1999) DNA strand breaks in Alzheimer’s disease. Brain Res 849, 67–77.

[5] Caldecott KW (2008) Single-strand break repair and genetic disease. Nat Rev Genet 9, 619–631.

[6] Suberbielle E, Djukic B, Evans M, Kim DH, Taneja P, Wang X, Finucane M, Knox J, Ho K, Devidze N, Masliah E, Mucke L (2015) DNA repair factor BRCA1 depletion occurs in Alzheimer brains and impairs cognitive function in mice. Nat Commun 6, 8897.

[7] McKinnon PJ (2017) Genome integrity and disease prevention in the nervous system. Genes Dev 31, 1180–1194.

[8] Shanbhag NM, Evans MD, Mao W, Nana AL, Seeley WW, Adame A, Rissman RA, Masliah E, Mucke L (2019) Early neuronal accumulation of DNA double strand breaks in Alzheimer’s disease. Acta Neuropathol Commun 7, 77.

[9] Thadathil N, Delotterie DF, Xiao J, Hori R, McDonald MP, Khan MM (2021) DNA Double-Strand Break Accumulation in Alzheimer’s Disease: Evidence from Experimental Models and Postmortem Human Brains. Mol Neurobiol 58, 118–131.

[10] Asada-Utsugi M, Uemura K, Ayaki T, M TU, Minamiyama S, Hikiami R, Morimura T, Shodai A, Ueki T, Takahashi R, Kinoshita A, Urushitani M (2022) Failure of DNA double-strand break repair by tau mediates Alzheimer’s disease pathology in vitro. Commun Biol 5, 358.

[11] Welch G, Tsai LH (2022) Mechanisms of DNA damage-mediated neurotoxicity in neurodegenerative disease. EMBO Rep 23, e54217.

[12] Dileep V, Boix CA, Mathys H, Marco A, Welch GM, Meharena HS, Loon A, Jeloka R, Peng Z, Bennett DA, Kellis M, Tsai LH (2023) Neuronal DNA double-strand breaks lead to genome structural variations and 3D genome disruption in neurodegeneration. Cell 186, 4404–4421 e4420.

[13] Suberbielle E, Sanchez PE, Kravitz AV, Wang X, Ho K, Eilertson K, Devidze N, Kreitzer AC, Mucke L (2013) Physiologic brain activity causes DNA double-strand breaks in neurons, with exacerbation by amyloid-beta. Nat Neurosci 16, 613–621.

[14] Weber Boutros S, Unni VK, Raber J (2022) An Adaptive Role for DNA Double-Strand Breaks in Hippocampus-Dependent Learning and Memory. Int J Mol Sci 23.

[15] Yap EL, Greenberg ME (2018) Activity-Regulated Transcription: Bridging the Gap between Neural Activity and Behavior. Neuron 100, 330–348.

[16] Madabhushi R, Gao F, Pfenning AR, Pan L, Yamakawa S, Seo J, Rueda R, Phan TX, Yamakawa H, Pao PC, Stott RT, Gjoneska E, Nott A, Cho S, Kellis M, Tsai LH (2015) Activity-Induced DNA Breaks Govern the Expression of Neuronal Early-Response Genes. Cell 161, 1592–1605.

[17] Delint-Ramirez I, Konada L, Heady L, Rueda R, Jacome ASV, Marlin E, Marchioni C, Segev A, Kritskiy O, Yamakawa S, Reiter AH, Tsai LH, Madabhushi R (2022) Calcineurin dephosphorylates topoisomerase IIbeta and regulates the formation of neuronal-activity-induced DNA breaks. Mol Cell 82, 3794–3809 e3798.

[18] Wu W, Hill SE, Nathan WJ, Paiano J, Callen E, Wang D, Shinoda K, van Wietmarschen N, Colon-Mercado JM, Zong D, De Pace R, Shih HY, Coon S, Parsadanian M, Pavani R, Hanzlikova H, Park S, Jung SK, McHugh PJ, Canela A, Chen C, Casellas R, Caldecott KW, Ward ME, Nussenzweig A (2021) Neuronal enhancers are hotspots for DNA single-strand break repair. Nature 593, 440–444.

[19] Reid DA, Reed PJ, Schlachetzki JCM, Nitulescu, II, Chou G, Tsui EC, Jones JR, Chandran S, Lu AT, McClain CA, Ooi JH, Wang TW, Lana AJ, Linker SB, Ricciardulli AS, Lau S, Schafer ST, Horvath S, Dixon JR, Hah N, Glass CK, Gage FH (2021) Incorporation of a nucleoside analog maps genome repair sites in postmitotic human neurons. Science 372, 91–94.

[20] Zhang X, Liu Y, Huang M, Gunewardena S, Haeri M, Swerdlow RH, Wang N (2023) Landscape of Double-Stranded DNA Breaks in Postmortem Brains from Alzheimer’s Disease and Non-Demented Individuals. J Alzheimers Dis 94, 519–535.

[21] Gupte R, Liu Z, Kraus WL (2017) PARPs and ADP-ribosylation: recent advances linking molecular functions to biological outcomes. Genes Dev 31, 101–126.

[22] Kim JA, Kruhlak M, Dotiwala F, Nussenzweig A, Haber JE (2007) Heterochromatin is refractory to gamma-H2AX modification in yeast and mammals. J Cell Biol 178, 209–218.

[23] McLean CY, Bristor D, Hiller M, Clarke SL, Schaar BT, Lowe CB, Wenger AM, Bejerano G (2010) GREAT improves functional interpretation of cis-regulatory regions. Nat Biotechnol 28, 495–501.

[24] Abarinov EV, Beaudin AE, Field MS, Perry CA, Allen RH, Stabler SP, Stover PJ (2013) Disruption of shmt1 impairs hippocampal neurogenesis and mnemonic function in mice. J Nutr 143, 1028–1035.

[25] Nativio R, Lan Y, Donahue G, Sidoli S, Berson A, Srinivasan AR, Shcherbakova O, Amlie-Wolf A, Nie J, Cui X, He C, Wang LS, Garcia BA, Trojanowski JQ, Bonini NM, Berger SL (2020) An integrated multi-omics approach identifies epigenetic alterations associated with Alzheimer’s disease. Nat Genet 52, 1024–1035.

[26] Allen M, Carrasquillo MM, Funk C, Heavner BD, Zou F, Younkin CS, Burgess JD, Chai H-S, Crook J, Eddy JA, Li H, Logsdon B, Peters MA, Dang KK, Wang X, Serie D, Wang C, Nguyen T, Lincoln S, Malphrus K, Bisceglio G, Li M, Golde TE, Mangravite LM, Asmann Y, Price ND, Petersen RC, Graff-Radford NR, Dickson DW, Younkin SG, Ertekin-Taner N (2016) Human whole genome genotype and transcriptome data for Alzheimer’s and other neurodegenerative diseases. Scientific Data 3, 160089.

[27] Mostafavi S, Gaiteri C, Sullivan SE, White CC, Tasaki S, Xu J, Taga M, Klein H-U, Patrick E, Komashko V, McCabe C, Smith R, Bradshaw EM, Root DE, Regev A, Yu L, Chibnik LB, Schneider JA, Young-Pearse TL, Bennett DA, De Jager PL (2018) A molecular network of the aging human brain provides insights into the pathology and cognitive decline of Alzheimer’s disease. Nature Neuroscience 21, 811–819.

[28] Zhang B, Gaiteri C, Bodea L-G, Wang Z, Mcelwee J, Podtelezhnikov A, Alexei, Zhang C, Xie T, Tran L, Dobrin R, Fluder E, Clurman B, Melquist S, Narayanan M, Suver C, Shah H, Mahajan M, Gillis T, Mysore J, Macdonald E, Marcy, Lamb R, John, Bennett A, David, Molony C, Stone J, David, Gudnason V, Myers J, Amanda, Schadt E, Eric, Neumann H, Zhu J, Emilsson V (2013) Integrated Systems Approach Identifies Genetic Nodes and Networks in Late-Onset Alzheimer’s Disease. Cell 153, 707–720.

[29] Morabito S, Miyoshi E, Michael N, Shahin S, Martini AC, Head E, Silva J, Leavy K, Perez-Rosendahl M, Swarup V (2021) Single-nucleus chromatin accessibility and transcriptomic characterization of Alzheimer’s disease. Nat Genet 53, 1143–1155.

[30] Vitelli V, Galbiati A, Iannelli F, Pessina F, Sharma S, d’Adda di Fagagna F (2017) Recent Advancements in DNA Damage-Transcription Crosstalk and High-Resolution Mapping of DNA Breaks. Annu Rev Genomics Hum Genet 18, 87–113.

[31] Ward IM, Chen J (2001) Histone H2AX is phosphorylated in an ATR-dependent manner in response to replicational stress. J Biol Chem 276, 47759–47762.

[32] Ward IM, Minn K, Chen J (2004) UV-induced ataxia-telangiectasia-mutated and Rad3-related (ATR) activation requires replication stress. J Biol Chem 279, 9677–9680.

[33] Reddy PH, Tripathi R, Troung Q, Tirumala K, Reddy TP, Anekonda V, Shirendeb UP, Calkins MJ, Reddy AP, Mao P, Manczak M (2012) Abnormal mitochondrial dynamics and synaptic degeneration as early events in Alzheimer’s disease: implications to mitochondria-targeted antioxidant therapeutics. Biochim Biophys Acta 1822, 639–649.

[34] Swerdlow RH (2023) The Alzheimer’s Disease Mitochondrial Cascade Hypothesis: A Current Overview. J Alzheimers Dis 92, 751–768.

[35] Ashleigh T, Swerdlow RH, Beal MF (2023) The role of mitochondrial dysfunction in Alzheimer’s disease pathogenesis. Alzheimers Dement 19, 333–342.

[36] Martinez-Garcia PM, Garcia-Torres M, Divina F, Terron-Bautista J, Delgado-Sainz I, Gomez-Vela F, Cortes-Ledesma F (2021) Genome-wide prediction of topoisomerase IIbeta binding by architectural factors and chromatin accessibility. PLoS Comput Biol 17, e1007814.

[37] Bramham CR, Worley PF, Moore MJ, Guzowski JF (2008) The immediate early gene arc/arg3.1: regulation, mechanisms, and function. J Neurosci 28, 11760–11767.

[38] Khazeem MM, Casement JW, Schlossmacher G, Kenneth NS, Sumbung NK, Chan JYT, McGow JF, Cowell IG, Austin CA (2022) TOP2B Is Required to Maintain the Adrenergic Neural Phenotype and for ATRA-Induced Differentiation of SH-SY5Y Neuroblastoma Cells. Mol Neurobiol 59, 5987–6008.

